# How prior pair-bonding experience affects future bonding behavior in monogamous prairie voles

**DOI:** 10.1101/2020.06.05.135160

**Authors:** Kelsey J. Harbert, Matteo Pellegrini, Katelyn M. Gordon, Zoe R. Donaldson

## Abstract

Monogamous prairie voles (*Microtus ochrogaster*) form mating-based pair bonds. Although wild prairie voles rarely re-pair following loss of a partner, laboratory studies have shown that previous pairing and mating does not negate the ability to form a new partner preference. However, little is known about how prior bond experience may alter the trajectory and display of a new pair bond. In the present study, we disrupted an initial pair bond by separating partners and then varied the amount of time before a new partner was introduced. We assessed how separation time affected the stability of partner preference over time and influenced decision-making in male voles performing a head-to-head partner preference test in which they chose between the first and second partner. We found that the ability to consistently display a preference for the second partner, supplanting the initial pair bond, depended on how long the test animal was separated from their first partner. Prior bonding experience also shaped the subsequent effects of mating on partner preference. Partner preference strength was sensitive to latency to mate with the second partner but not the first partner, irrespective of separation time. These results suggest that the ability to form a consistent, strong preference for a new partner after an initial pair bond depends upon the amount of time that has passed since separation from the first partner. These results provide valuable insight into how social bonds are dynamically shaped by prior social experience and identify variables that contribute to recovery from partner loss and the ability to form a new pair bond.

---

Romantic relationships are dynamic over time. It is not uncommon for humans to sequentially form more than one pair bond. The formation and dissolution of these bonds can have profound effects on emotional well-being and health (Holt-Lunstad et al., 2010; Keyes et al., 2014; Shor et al., 2012). Given the importance of social bonds, understanding how previous relationships and other experiential factors impact subsequent bonds has the potential to elucidate the complex interplay between bonding and health.

Monogamous prairie voles (*Microtus ochrogaster*) form exclusive pair bonds, providing a tractable laboratory species for exploring the factors that contribute to the ability to form a new bond in bond-experienced individuals. In the wild, prairie voles occasionally re-pair; ~20% will take a new partner (Carter and Getz, 1993). Recent laboratory studies suggest that most male voles will form a partner preference, a behavioral proxy for a pair bond, regardless of prior pairing, even after being sequentially paired and mating with up to 10 females (Kenkel et al., 2019). While this demonstrates that previous pairing/mating does not negate the ability to form future bonds, a number of questions remain regarding the role of previous experience in subsequent bond formation and expression. For instance, it remains unknown how the behavioral characteristics of the second bond compares to the first bond and how time between separation from the first partner and introduction to the second partner affects partner preference.

To further define the effects of previous pair bond experience on the subsequent formation and stability of a new bond, we performed a controlled experiment of sequential pairings of male prairie voles with female partners. The presence, strength, and consistency of each pair bond were measured using multiple partner preference tests (PPTs). This assay tracks how much time the test animal spends with their partner versus a novel, opposite-sex vole tethered at opposite ends of a testing apparatus (Williams et al., 1992). By varying time between removal of the first partner and introduction to the second partner, we found that 4 weeks of separation is required for the formation of a stable second pair bond, and likewise, that only after 4 weeks was the first bond supplanted by the second in a head-to-head test. In addition, we found an effect of mating latency on partner preference only with second partners, suggesting a role for previous pairing/mating experience in shaping subsequent bonds. Together, this indicates that not all bonds are the same and provides insight into variables that contribute to the ability to form a new bond following the loss of a prior bond.

## Methods

### Animals

Sexually naive adult prairie voles (N = 66: 22M, 44F) were bred in-house in a colony originating from a cross between voles obtained from colonies at Emory University and University of California Davis, both of which were established from wild animals collected in Illinois. Animals were weaned at 21 days and housed in same-sex groups of 2 – 4 animals in standard static rodent cages (7.5 × 11.75 × 5 in.) with ad-lib water, rabbit chow (5326-3 by PMI Lab Diet) supplemented with alfalfa cubes, sunflower seeds, cotton nestlets, and igloos for enrichment until initiation of the experiment. In order to eliminate confounds of pregnancy, females were tubally ligated and given at least two weeks to recover prior to the start of the experiment (details below). All voles were between the ages of 8 and 16 weeks at the start of the experiment. Throughout the experiment, animals were housed in smaller static rodent cages (11.0 in. × 8.0 in. × 6.5 in.) with ad-lib water, rabbit chow, and cotton nestlets. They were kept at 23–26°C with a 10:14 dark: light cycle. All procedures were approved by the University of Colorado Institutional Animal Care and Use Committee.

### Tubal ligation

Tubal ligation surgeries were performed using a dorsal approach and isoflurane as an anesthetic (Souza et al., 2019). An electric razor was used to clear the immediate area of fur, and iodine was applied to the disinfect the exposed skin. A central vertical cut was made through the skin just below the ribs. The opening was pulled over to one side, and an internal incision was made through the abdominal wall above the ovary. The ovary and upper uterus were briefly removed from that abdomen, and a cauterizing tool was used to simultaneously bisect and seal the edges of the upper uterus while leaving the ovary untouched. The tissues were replaced in the body cavity. The internal incision was sutured using vicryl-coated sutures, size 4-0, and the procedure was repeated on the other side. The external incision was then sealed with surgical staples, which were removed one-week post-surgery. All females were given at least 2 weeks to recover prior to their first pairing. Efficacy of tubal ligation was demonstrated by the fact that none of our females became pregnant despite visual evidence of mating.

### Experimental Design

All test males were paired with a female partner (Partner 1) on opposite sides of a custom transparent, ventilated, divider, which reliably induces sexual receptivity in the female voles and decreases aggression (Donaldson et al., 2009). Dividers were removed after 48 hours and sexual behavior was recorded from the side of the cage via Sony Handycams (DCR-SX85) with four cages captured per frame, for the first 3 hours following divider removal. PPTs (procedure described below) were performed at short-term (3 days post-divider removal) and long-term (10 days post-divider removal) timepoints, enabling us to investigate changes in pair bond strength as a function of pairing time (Scribner et al., 2020). For all PPT tests, except for the final head-to-head test, novel females consisted of partners from other pairings. Test animals were never re-exposed to the same novel animals nor were they paired with or exposed to a sibling during PPT. Immediately following the long-term PPT, the partners were separated and singly housed for 48 hours, 2 weeks, or 4 weeks, randomly assigned. After the isolation period, pairing and PPT testing were repeated with a new sexually naive partner (Partner 2) following the same timeline as for Partner 1. Following the long-term PPT with Partner 2, all voles were singly housed for 48 hours. A final PPT was performed in which the test animal chose between Partner 1 and Partner 2 tethered in opposite chambers. This final head-to-head PPT was designed to determine whether the first or second bond was behaviorally dominant (Fig. 1).

**Figure 1.**
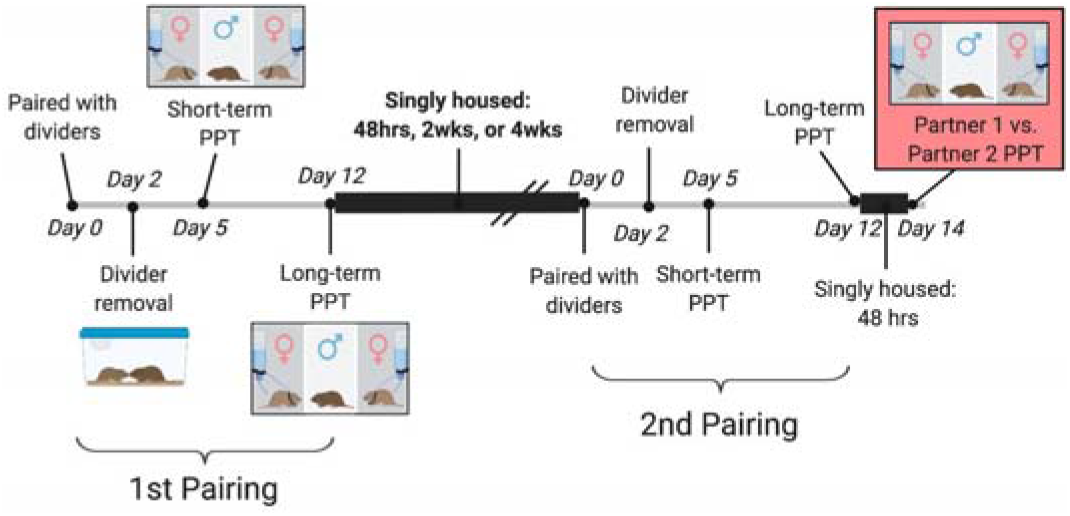
Experiment Design. Male voles were paired with naïve females and placed in small cages with a lengthwise divider for 2 days. Once dividers were removed, mating was recorded for 3 hours, and then continued to live with their opposite-sex partner. 3 days after divider removal, males underwent a short-term PPT. The pairs then cohabitated for another 7 days before undergoing a long-term PPT. Immediately following the long-term PPT, all voles were singly housed according to their assigned separation condition of 48 hours, 2 weeks, or 4 weeks. Test animals were then paired with a new sexually-naïve female in a divided cage and underwent the same testing schedule as in their first pairing. Immediately after the long-term PPT with partner 2, all voles were singly housed for 2 days at which point the final, head-to-head PPT was performed. In this PPT the males chose between tethered Partner 1 and Partner 2 to determine whether the new bond formed with Partner 2 supplanted the bond formed with Partner 1.

### Partner Preference Test

Each PPT apparatus consisted of a box (75.0 cm. long × 20.0 cm. wide × 30.0 cm. tall) sectioned into three equal size chambers separated by removable dividers (Scribner et al., 2020). Testing was carried out as described in Ahern et al., (2009). Partner and a novel age-matched conspecifics were tethered to bolts located on opposite sides of the apparatus using fishing swivels and zip ties with a water bottle affixed to the same wall. Two alfalfa pellets were placed in each chamber containing a tethered animal. Overhead cameras (Panasonic WVCP304) were used to film two boxes simultaneously. The test animal was placed in the middle chamber, dividers were removed, and it freely explored the apparatus for 3 hours. At the end of the test, the apparatus was cleaned, and a second test was performed; the partner for the first test animal served as the novel for the second and vice versa. The movement of the test animal was recorded and tracked post-hoc using Topscan High-Throughput v3.0 software (Cleversys Inc.) using the parameters from Ahern et al. (2009). Frame by frame behavioral data was analyzed using a custom Matlab script to calculate the average distance between the test animal and tethered animal when in the same chamber, time spent huddling with each tethered animal, and total distance traveled. The partner preference score was calculated using Partner Huddle/Partner + Novel Huddle.

### Statistics

Data were analyzed using SPSS version 25. Details of all statistical tests are provided in Supplementary Table 1. As a behavioral test, comparison of time spent with the partner versus the novel animal violates the assumptions of a traditional T-test because time with each tethered vole is not truly independent. To address this, partner preference was assessed using the preference score. This was compared to a null value of 0.5 (no preference) in a two-tailed one-sample T-test. Differences in preference across conditions and/or testing timepoints were analyzed using an RM-ANOVA with Timepoint as a within-subject factor and Condition as a between-subject factor. To gain further insight into the underlying behavioral changes that contributed to differences in partner preference scores over time, total partner huddle or total novel huddle across timepoints were analyzed separately using a paired T-test. To determine behavioral consistency across timepoints, we examined correlations between the total partner huddle time, novel huddle time, and preference score between short-term and long-term timepoints.

To strengthen our interpretation, we also examined the average distance between the test animal and the tethered animals when the test animal was in the chamber with the tethered animal. We have previously shown that this behavioral metric serves as a proxy for partner preference (Scribner et al., 2020). Because the distance from the partner while in the partner chamber does not influence the distance from the novel in the novel chamber, these variables can be considered independent, and we performed a paired T-test and/or RM-ANOVA to determine whether these metrics differed within and across tests. In addition, we calculated a ratio of novel distance:partner distance to create a within-animal preference score based on distance and asked whether this score correlated with the preference score.

Finally, we examined the effects of mating latency on partner preference. We performed a Kaplan Meyer survival analysis with Log Rank for overall comparison to examine potential group differences in mating latency. This approach provides an ideal non-parametric test that takes into account failure to complete the task (e.g. failure to mate).

## Results

### Excluded Animals

Eight animals (out of an original 66) were excluded from one or more PPT due to unanticipated partner losses or technical problems (aggression towards the stranger in the partner preference test: n = 5, escape from the partner preference test: n = 1, or a faulty camera attachment: n = 2). The PPT from which each animal was excluded are listed in Supplemental Table 2, and their exclusion of different statistical tests is listed in Supplemental Table 1.

### Preference for Partner 1 is evident at short-term and long-term timepoints

#### Partner preference

We measured partner preference at two timepoints to assess potential changes in bond-related behaviors as a function of time paired. Relative to a null hypothesis of no preference, sexually naive male voles paired with a female partner demonstrated a partner preference at short-term (p = 0.017) and long-term (p < 0.001) timepoints (Fig. 2A). There was no significant change in preference scores between short-term and long-term tests (p = 0.197), so we next examined whether there were changes in either partner or novel huddle time, respectively, across timepoints. Time spent with the partner did not change over time (p = 0.849), but we did observe a decrease in novel huddle time, suggesting a strengthening of partner preference through decreased novel interaction (Fig. 2B; p = 0.025). We also examined the average distance between the test animal and the tethered animal while they were in the same chamber. At both timepoints, test animals were physically closer to their partner than the novel animal when in the chamber with them (Fig. 2C; main effect of Tethered animal; p < 0.001). These two metrics – preference score and average distance ration (P/N) – were correlated strongly at both testing timepoints (Fig. 2D; short-term: R = 0.78, p < 0.001; long-term: R = 0.76, p < 0.001), indicating that they measured overlapping aspects of preference behavior and can both be used as proxies for inferring partner preference.

**Figure 2.**
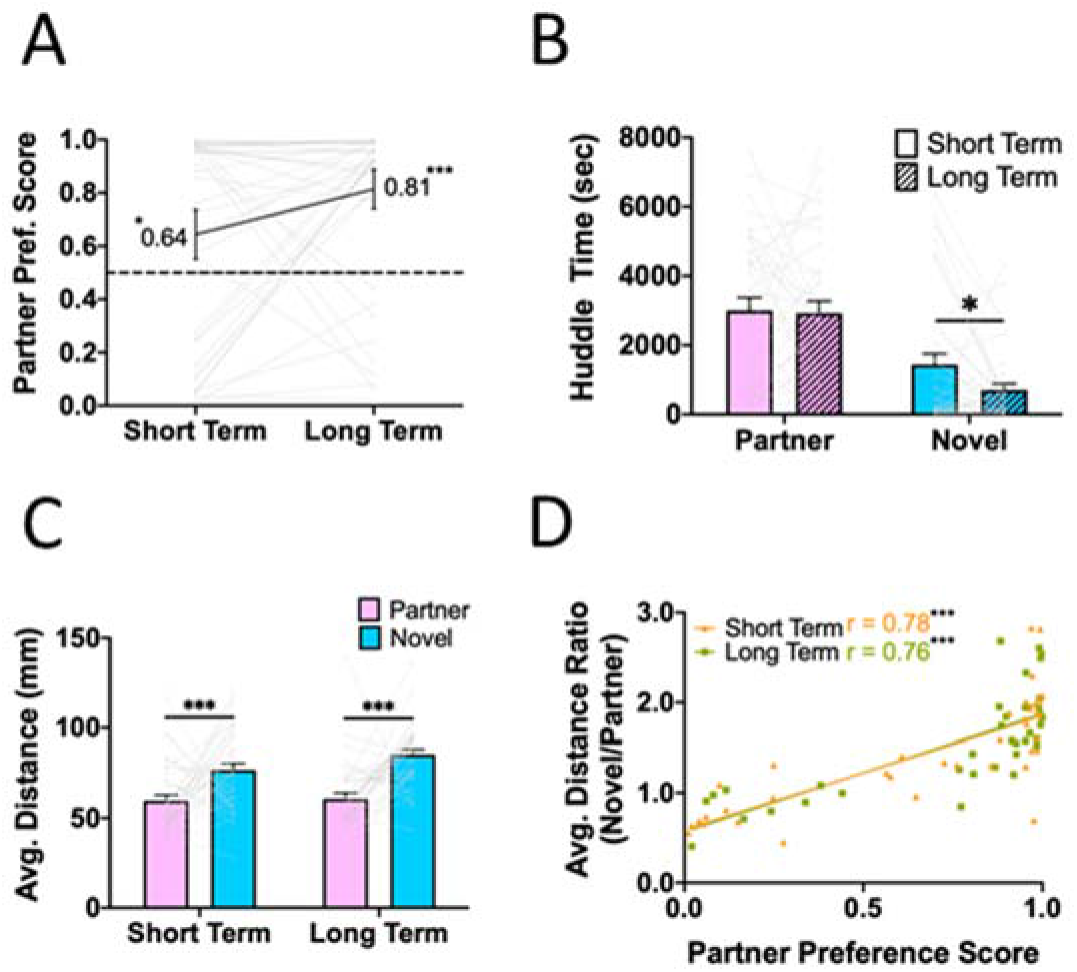
Metrics of partner preference for Partner 1. Male prairie voles housed with a sexually receptive tubally-ligated female prairie vole exhibited a partner preference following 3 days (short-term) and 10 days (long-term) of cohabitation post-divider removal. **A)** Partner preference score (proportion of time spent huddling with the partner) was significantly greater than chance (0.5) at both timepoints (short-term: p = 0.017, long-term: p < 0.001). **B)** Time spent huddling with the partner did not change over time (p = 0.849), while time spent huddling with the novel decreased over time (p = 0.025). **C)** Average distance from the tethered animal while in the same chamber also reflects partner preference. At both timepoints, the test animal was physically closer to their partner when in the partner chamber than they were to the novel animal when they were in the novel chamber (main effect of tethered animal: p < 0.001). **D)** To examine the consistency between preference score and distance metrics, we calculated a distance ratio (novel distance/partner distance) from (C). There was a strong correlation between the preference score and the distance ratio at both timepoints (short-term: p < 0.001; long-term: p < 0.001), suggesting that both metrics provide valid estimates of partner preference. Significance notated as: *p < 0.05, **p < 0.005, ***p < 0.0005.

#### Behavioral consistency

Time spent huddling with the partner was positively correlated across the short-term and long-term tests, with a similar trend for novel huddle time (Fig. 2D partner: R = 0.347, p = 0.036; novel: R = 0.294, p = 0.077), suggesting at least moderate intra-animal behavioral consistency across tests. Similarly, the total distance traveled within the test chamber was also positively correlated (R = 0.370; p = 0.026), although there was a decrease in total locomotion between the short-term and long-term test, suggesting potential habituation to the testing environment (main effect of Timepoint: p = 0.001).

### Stability for preference for Partner 2 depends on separation time

Male voles were randomly assigned to different separation times (48 Hour, n = 15; 2 Week, n = 11; 4 Week, n = 11). The conditions did not differ in initial preference for Partner 1, total distance traveled in PPTs with the first partner, or mating latency. Detailed statistical comparisons are available in Supplemental Table 1. Thus, these conditions were behaviorally equivalent with respect to the behaviors displayed towards their first partner. Similar to previous reports by Kenkel et al. (2019), we found that voles in all conditions were capable of showing a preference for their second partner within 3 days of pairing (short-term; 48 Hour p = 0.022, 2 Week p = 0.003, 4 Week p = 0.075 (but see significant distance data), Fig. 3: A, D, G). However, when re-tested following a longer cohabitation, only animals in the 4 Week condition showed a significant partner preference at the long-term timepoint (48 Hour p = 0.285, 2 Week p = 0.850, 4 Week p = 0.016, Fig. 3: A, D, G). These results were consistent with those observed for average distance from the tethered animal. Specifically, males from the 48 Hour and 2 Week conditions were closer to the partner than the novel animals at the short-term timepoint, but did not show a significant difference at the long-term timepoint (48 Hour: short-term p = 0.285 long-term p = 0.196; 2 Week: short-term p = 0.005 long-term p = 0.549; Fig. 3: C, F). In contrast, the 4 Week condition was closer to the partner than the novel at both timepoints (4 Week: short-term p = 0.042 long-term p = 0.002; Fig 3I). As with the first pairing, partner huddle and novel huddle were positively correlated across tests for Partner 2 (all conditions combined; partner: R = 0.660, p < 0.001, novel: R = 0.614, p < 0.001). Total distance traveled in the test apparatus was not correlated across tests (R = 0.215, p = 0.229), although the animals in the 4 Week condition showed consistently lower levels of locomotion than the other two conditions (main effect of Condition: p = 0.0004). Together, this suggests a potential rebound effect in which all animals initially show a preference for their second partner, but this remains consistent over time only for males in the 4 Week condition.

**Figure 3.**
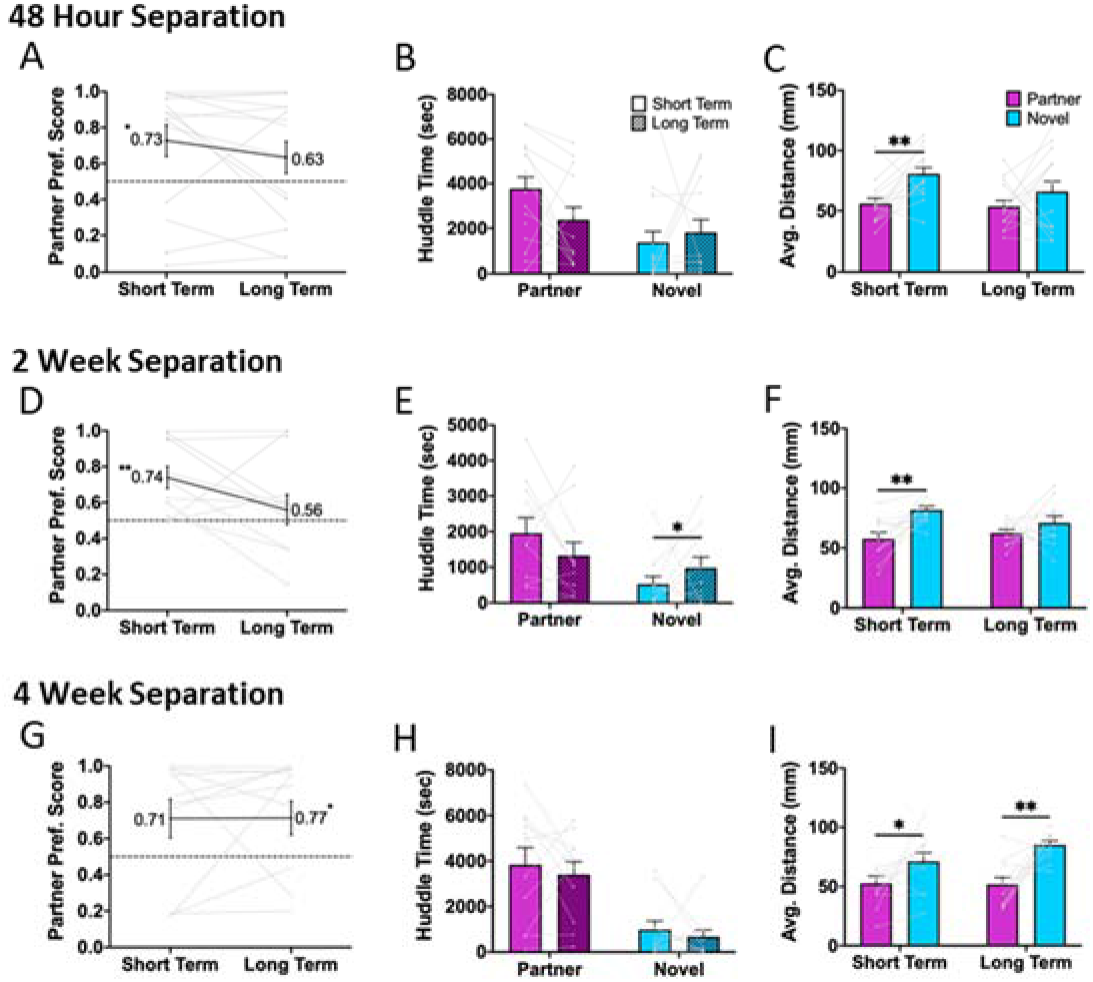
Metrics of partner preference for Partner 2. Male prairie voles spent 48 hours, 2 weeks, or 4 weeks singly housed between separation from their first partner and introduction to their second partner. As with Partner 1, animals were tested for partner preference at 3 days (short-term) and 10 days (long-term) post cage divider removal. *48 Hour Separation:* **A)** Male voles showed a partner preference at the short-term test (p = 0.022) but not at the long-term test (p = 0.285). **B)** There were no changes in partner huddle time (p = 0.092) across tests, nor in novel huddle time (p = 0.172) across tests. **C)** Males were closer to their partner than the novel animal during the short-term (p = 0.002) but not the long-term test (p = 0.196). *2 Week Separation:* **D)** Partner preference was evident at the short-term (p = 0.003) but not the long-term (p = 0.850) test. **E)** Time spent huddling with the partner did not change between tests (p = 0.131) while time spent huddling with the novel increased (p = 0.032). **F)** Males were closer with their partner than the novel at the short-term test (p = 0.005) but not at the long-term test (p = 0.549). *4 Week Separation:* G) there was a trend towards partner preference at the short-term timepoint (p = 0.075), that was fully evident at the long-term (0.016). **H)** Neither time spent huddling with the partner (p = 0.577) nor time spent huddling with the novel (p = 0.571) varied over time. **I)** Test animals were closer in proximity to the partner than the novel at both the short-term (p = 0.042) and long-term (p = 0.002) tests. Notably, males in the 4 week separation group were the only males to show a partner preference at the long-term test, which is reflected in a consistent decreased distance from the partner compared with the novel.

### Mating latency predicts partner preference for Partner 2 but not for Partner 1

Previous work suggests that mating facilitates partner preference. Thus, we separated animals into early and non-early mating groups based on whether they mated within 3 hours of divider removal. There were no significant differences in mating latency across pairings or between conditions, indicating that likelihood to mate within the first 3 hours after divider removal is not influenced by prior pairing or by separation time. There were no differences between the first and second pairings (p = 0.452; Fig. 4C). In both instances, a similar proportion of animals (62% in the first pairing and 49% in the second pairing) mated within the first 180 minutes (p = 0.187; Fig. 4A, B). When analyzed by separation condition (48 Hour, 2 Week, 4 Week), there was no significant difference in latency to mate between conditions for either pairing (first pairing p = 0.505, second pairing p = 0.653) (Fig. S1A, B).

**Figure 4.**
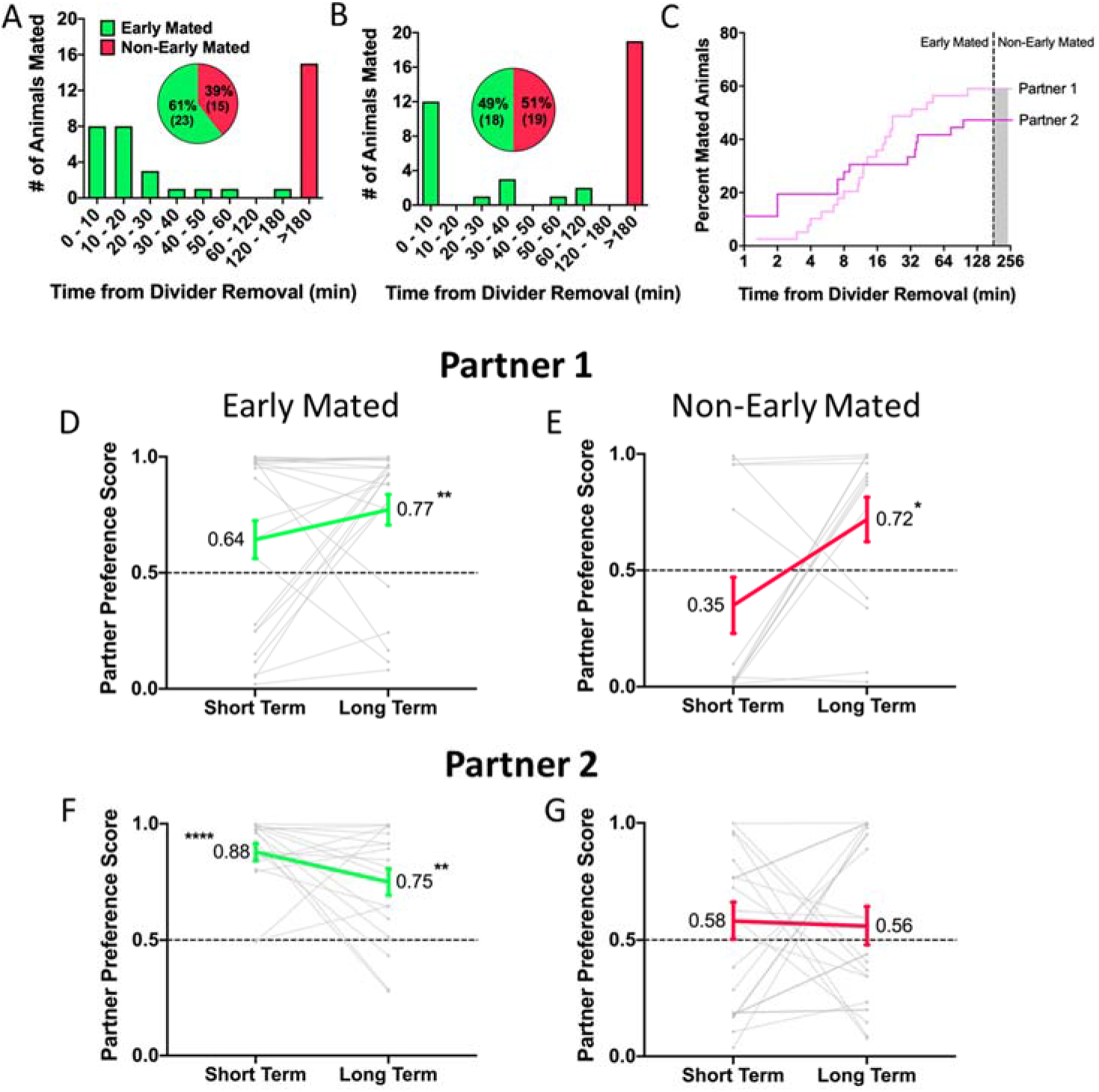
Effects of mating latency on partner preference. Dividers were removed from cages 48 hours post-pairing, and mating behavior was filmed and scored for the first 180 minutes of interactions. Voles that mated within the first 180 minutes were classified as “Early Mated” while those who did not were classified as “Non-Early Mated”. A, B) For the first and second pairings, approximately half of the test animals mated within the first 180 minutes, and there were no differences in the proportion of early mated animals for partner 1 vs partner 2 (p = 0.187). C) Likewise, there were no significant differences in latency to mate with Partner 1 or Partner 2 (p = 0.452). D, E) Partner preference emerged at the long-term PPT for both early (D) and non-early (E) maters (Early mated: short term, p = 0.094, long term p = 0.0004;. Non-early mated: short term, p = 0.099, long term p = 0.039) F) Only animals that mated within the first 180 minutes showed a preference for partner 2 (short term, p < 0.001, long term, p = 0.002). G) Non-early maters failed to form a partner preference at either time point (short term, p = 0.309, long term, p = 0.623).

We next examined whether mating latency predicted differences in bond strength (preference score) for either partner. There were no differences in preference for Partner 1 between early and non-early mated males (main effect of Latency: p = 0.279) (Fig. 4D, E). In contrast, mating latency strongly predicted preference for Partner 2, with non-early maters failing to show a partner preference (main effect of Latency, p = 0.005) (Fig. 4F, G). This suggests that prior pairing leads to a stronger effect of mating latency on subsequent preference formation.

### Four weeks of separation are required to supplant an old bond with a new one

Finally, we asked whether preference for Partner 1 or Partner 2 predominated in a head-to-head PPT. There were no consistent preferences for Partner 1 vs. Partner 2 for animals in the 48 Hour and 2 Week separation conditions (48 Hour: p = 0.981, 2 Week: p = 0.406; Fig. 5A). However, males in the 4 Week condition consistently spent more time huddling with Partner 2 than Partner 1 (p < 0.001; Fig. 5A). This was similarly evident when we examined average distance from the tethered animals when the test animal was in the chamber (48 Hour p = 0.573; 2 Week p = 0.315; 4 Week p < 0.001) (Fig. 5B). There were no differences in total locomotion during the head-to-head PPT across separation conditions (main effect of Condition: p = 0.998). This suggests that 4 weeks of separation prior to the introduction of a new partner leads to a pair bond that supplants the prior bond, but this does not occur following shorter periods of separation.

**Figure 5.**
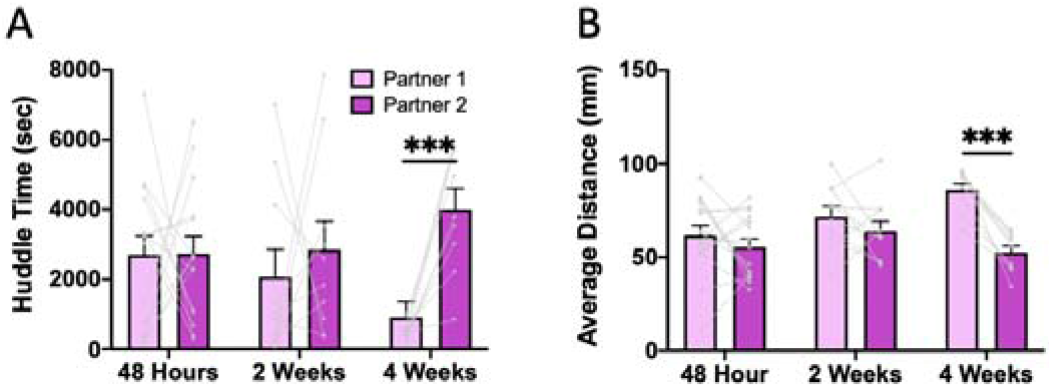
Metrics for the head-to-head partner preference test. Voles were singly housed for 48 hours following the long-term PPT with Partner 2. Test animals were then placed in a PPT with Partner 1 and Partner 2 as the tethered animals. **A)** Males in the 4 Week group were the only test animals to spend significantly more time huddling (p < 0.001) with Partner 2 than Partner 1 (48 hour, p = 0.981; 2 Week, p = 0.406). **B)** Test animals in the 4 Week group stayed significantly closer (p < 0.001) to Partner 2 than Partner 1.While test animals in the 48 Hour (p = 0.573) and 2 Week (p = 0.315) groups did not show a difference in proximity to Partner 1 versus Partner 2.

## DISCUSSION

The goal of this study was to examine how a previous pair bond altered subsequent bonding behavior with a new partner. We found that male prairie voles formed a new partner preference following the loss of their first partner. However, the formation and stability of that preference, as well as whether the new preference supplants the old one, depend on mating latency and separation time. Only voles separated from the first partner for 4 weeks formed a consistent second bond that supplanted the first. Overall this suggests that the full dissolution of a pair bond as measured by partner preference is time-dependent. Further, while mating latency does not predict the strength of a vole’s first bond, once an animal has experienced a bond, mating latency becomes a more important predictor of successful re-bonding. This experience-dependent effect suggests that voles may apply previously learned information about factors that affect bond success. Together, this indicates that subsequent pair bonds are shaped by initial bonding experience and subsequent separation time.

A prior study opportunistically used “stud males,” or male voles that were known to reliably mate, to show that that male prairie voles can demonstrate a partner preference even following pairing and mating with up to 10 females (Kenkel et al., 2019). Our study builds on this initial observation in three key ways. As discussed in more detail below, we measured latency to mate, stability of preference for the same partner over time, and finally, we used a head-to-head test to determine the primacy of the first versus second bond. Incorporating these metrics provides an additional layer to our understanding of the role of previous social experience on future attachment formation.

We found that longer latency to mate systematically predicts weaker partner preference only for the second partner and not the first. This was somewhat surprising given that previous work has shown that mating can also affect initial partner preference (Williams, 1992; Williams et al., 1992). However, in the previous study, they measured partner preference 6 hours after pairing. We initially assessed partner preference 3 days after pairing and 5 days after introduction via dividers. It is possible that mating latency has a stronger effect on preference for a first partner during early pairing/cohabitation, which is supported by our observation that the non-early maters exhibited, if anything, a nominal novel preference at the short-term timepoint prior to forming a preference at the long-term timepoint. This could be an effect of experiential familiarity, meaning, once animals are no longer sexually naive, the propensity to mate soon after divider removal may play a larger role in their assessment of bond quality.

We also examined the role of cohabitation time on preference strength. This enabled us to determine whether preferences change over time. We have previously reported that, using an abbreviated, 20-minute PPT, prairie voles demonstrate a strengthening of preference over time. Here we detected a trend for stronger partner preference for the first partner at the long-term timepoint, consistent with our previous result, although this did not reach statistical significance. Strikingly, while this trend was also observed for the second partner in the 4 Week separation condition, voles with a shorter separation duration exhibited a *decrease* in partner preference at the long-term timepoint. The fact that this occurred in both the 48 Hour and 2 Week separation conditions suggests that preference reliably decreases if insufficient time has passed before pairing with a new partner. However, it is worth noting that almost all voles showed a preference for one partner, and the condition-level data do not reflect individual lack of preference. Together with the results of our head-to-head test, as detailed below, this may suggest that the initial pair bond is not yet fully dissolved.

Finally, we hypothesized that separation time would predict whether the test animal preferred their first or second partner in a head-to-head test. This appears to be partially true; only voles separated from their first partner for at least 4 weeks reliably choose their second partner in a head-to-head test. However, our shortest separation timepoint did not reliably result in a preference for the first partner, suggesting that while there is an upper limit on how long a previous pair bond can predominate, there is also tremendous individual variation in how quickly a second bond can supplant the first. This finding is parsimonious with a previous study that showed that prairie voles no longer show a preference for their partner after four weeks of separation (Sun et al., 2014). However, of note, isolation may be key for the dissolution of partner preference, as animals in the 2 Week condition had also not seen their initial partner in 4 weeks at the time of the head-to-head test, yet a subset still preferred their initial partner.

Partner preference is an inherently complex task dependent on internal state, as well as ongoing social interactions that occur between the test and tethered animals throughout the 3-hour test. Accordingly, there is marked variation in partner preference across multiple trials such that some animals even show a decrease in preference for their first partner at the long-term timepoint. A previous study suggested that the amount of time spent huddling with different interaction partner (same-sex, opposite-sex, familiar vs. unfamiliar) was not consistent for a given vole (Ahern et al., 2019). Thus, we asked whether voles exhibit consistency in huddling behavior for the same partner in the same task. We found that for both the first and second partner, the amount of time spent huddling with the partner or with the novel was correlated across the short-term and long-term tests. The preference score was also correlated, albeit more weakly. This suggests that prairie voles are consistent in their huddling behavior when engaging in equivalent scenarios, such as sequential PPTs for the same partner. In contrast, the previously noted differences in huddling during different social interactions may reflect a level of social decision-making that incorporates differences in social valence or other factors related to interaction with different individuals (Ahern et al., 2019).

It is worth noting that at a superficial level, our results and those of Kenkel et al (2019), which indicate that preference formation is common even following multiple pairings, may seem at odds with reports that only ~20% of male and female prairie voles re-pair in the wild (Carter and Getz, 1993). On the contrary, our results, which indicate that 4 weeks of separation from the first partner is required in order to form a new bond that supplants the old one, support the low rates of re-pairing in the wild. In particular, the average life expectancy of prairie voles in the wild is 65.6 +/− 1.7 days (Getz et al., 1997). If an animal pairs and subsequently loses a partner, it is quite possible that they will not survive the 4 weeks that appear to be needed to overcome that initial bond and form a new one. In addition, rebonding requires the availability of a non-bonded opposite-sex animal, which in some populations may represent a limiting factor (Getz et al., 1997). Finally, work with semi-natural vole populations suggests that aggression towards likely partners contributes to a failure to rebond (Thomas and Wolff, 2004). It is not clear how long this aggression typically lasts in the wild, but in our study, we used cage dividers to habituate test males to their partners, which reduced aggression, induced behavioral receptivity in the females, and enabled introduction of a new partner in as few as 48 hours after removal of the first partner. Thus, while our experimental design was optimized to ask whether and how strongly voles rebond following separation from a first partner, extrapolation of our results suggests that the conditions needed for rebonding are rarely met in wild populations.

Finally, our study has a few limitations that represent areas of future inquiry. Specifically, our study was limited to males, and we do not have data on multiple bond formation in females. The reproductive demands for females differ dramatically from those of males, and as such, there may be sexual dimorphism and/or a role for pregnancy in the propensity to rebond. In addition, it remains unclear what the mechanisms contribute to the predominance of the second bond in the head-to-head tests of our 4 Week separation condition. This could be due to forgetting the first partner or failing to associate the motivational salience with that individual independent of recognition, e.g. bond dissolution. Thus future studies are needed to dissociate these two mechanisms.

In sum, the present study provides a foundation upon which we can investigate the neurobiological mechanisms subserving adaptation to bond dissolution in a species whose social biology resembles that of humans. Humans often form more than one pair bond, but the mechanisms that contribute to adaptation to partner loss and enable a new bond to form remain largely unexplored. By showing that male voles can form stable second bonds when given adequate time following separation from their first partner, we have provided the beginnings of a behavioral model for studying adaptation to loss and subsequent rebonding that might one day be translated to clinical interventions for humans struggling to overcome partner loss.

## Author Contributions and Notes

Z.R.D., H.K., and M.P designed research, H.K., M.P. and K.G. performed research, Z.R.D. analyzed data; and H.K. and Z.R.D. wrote the paper.

The authors declare no conflict of interest.

This article contains supporting information online.

## Acknowledgments

We would like to thank the animal care staff at University of Colorado Boulder. This work was supported by awards from the Dana Foundation, the Whitehall Foundation, and NIH DP2OD026143 to ZRD.

## References

Ahern, T.H., Modi, M.E., Burkett, J.P., Young, L.J., 2009. Evaluation of two automated metrics for analyzing partner preference tests. Journal of neuroscience methods 182, 180–8. https://doi.org/10.1016/j.jneumeth.2009.06.010

Carter, C.S., Getz, L.L., 1993. Monogamy and the prairie vole. Sci. Am 268, 100–106.

Donaldson, Z.R., Yang, S.-H., Chan, A.W.S., Young, L.J., 2009. Production of Germline Transgenic Prairie Voles (Microtus ochrogaster) Using Lentiviral Vectors1. Biology of Reproduction 81, 1189–1195. https://doi.org/10.1095/biolreprod.109.077529

Getz, L.L., Simms, L.E., McGuire, B., Snarski, M.E., 1997. Factors Affecting Life Expectancy of the Prairie Vole, Microtus ochrogaster. Oikos 80, 362. https://doi.org/10.2307/3546604

Holt-Lunstad, J., Smith, T.B., Layton, J.B., 2010. Social Relationships and Mortality Risk: A Meta-analytic Review. PLOS Medicine 7, e1000316. https://doi.org/10.1371/journal.pmed.1000316

Kenkel, W.M., Perkeybile, A.M., Yee, J.R., Carter, C.S., 2019. Rewritable fidelity: How repeated pairings and age influence subsequent pair-bond formation in male prairie voles. Hormones and Behavior 113, 47–54. https://doi.org/10.1016/j.yhbeh.2019.04.015

Keyes, K.M., Pratt, C., Galea, S., McLaughlin, K.A., Koenen, K.C., Shear, M.K., 2014. The burden of loss: unexpected death of a loved one and psychiatric disorders across the life course in a national study. Am J Psychiatry 171, 864–71. https://doi.org/10.1176/appi.ajp.2014.13081132

Scribner, J.L., Vance, E.A., Protter, D.S.W., Sheeran, W.M., Saslow, E., Cameron, R.T., Klein, E.M., Jimenez, J.C., Kheirbek, M.A., Donaldson, Z.R., 2020. A neuronal signature for monogamous reunion. Proceedings of the National Academy of Sciences 117, 11076–11084. https://doi.org/10.1073/pnas.1917287117

Shor, E., Roelfs, D.J., Curreli, M., Clemow, L., Burg, M.M., Schwartz, J.E., 2012. Widowhood and Mortality: A Meta-Analysis and Meta-Regression. Demography 49, 575–606. https://doi.org/10.1007/s13524-012-0096-x

Souza, V.R., Mendes, E., Casaro, M., Antiorio, A.T.F.B., Oliveira, F.A., Ferreira, C.M., 2019. Description of Ovariectomy Protocol in Mice. Methods Mol. Biol. 1916, 303–309. https://doi.org/10.1007/978-1-4939-8994-2_29

Sun, P., Smith, A.S., Lei, K., Liu, Y., Wang, Z., 2014. Breaking bonds in male prairie vole: Long-term effects on emotional and social behavior, physiology, and neurochemistry. Behavioural Brain Research 265, 22–31. https://doi.org/10.1016/j.bbr.2014.02.016

Thomas, S.A., Wolff, J.O., 2004. Pair bonding and “the widow effect” in female prairie voles. Behav. Processes 67, 47–54. https://doi.org/10.1016/j.beproc.2004.02.004

Williams, J., 1992. Development of partner preferences in female prairie voles (Microtus ochrogaster): The role of social and sexual experience. Hormones and Behavior 26, 339–349. https://doi.org/10.1016/0018-506X(92)90004-F

Williams, J.R., Carter, C.S., Insel, T., 1992. Partner preference development in female prairie voles is facilitated by mating or the central infusion of oxytocin. Annals of the New York Academy of Sciences 652, 487–9.

