## Supplemental tables and figure for "How prior pair-bonding experience affects future bonding behavior in monogamous prairie voles"

### Supplemental Materials

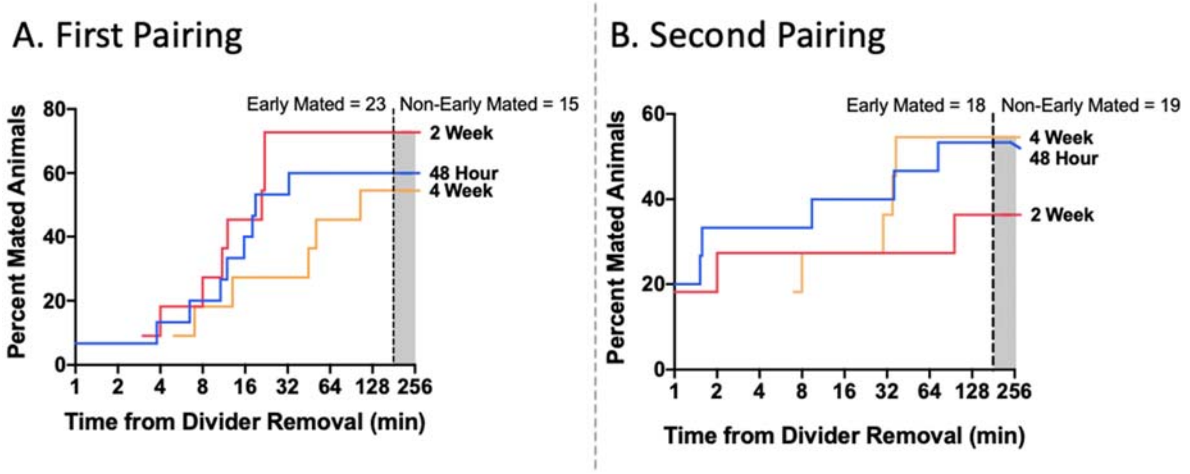

**Supplemental Figure 1. Mating latency comparison across 3 separation groups. A)** There were no significant group differences in latency to mate with the first partner (p = 0.505). **B)** There were no significant group differences in latency to mate with the second partner (p = 0.653).

| Experiment | Partner | Measurement | Statistical Test | Comparison | ° of freedom, error | F or T | p | * | Group Size | Fig. | Notes |
|---|---|---|---|---|---|---|---|---|---|---|---|
| Partner preference | 1 | Prop. Partner huddle | RM-ANOVA | Time (RM) | 1, 36 | 1.726 | 0.197 | | n = 37 | 1A | no data for 2242, long term PPT |
| | | Short Term | one way T-test | relative to 50% | 37 | 2.495 | 0.017 | * | n = 38 | | |
| | | Long Term | one way T-test | relative to 50% | 36 | 4.667 | 0.000 | *** | n = 37 | | |
| | | Group differences prop. Partner huddle | RM-ANOVA | Time (RM) | 1, 34 | 1.444 | 0.238 | | 48 hr = 15, 2 wk = 11, 4 wk = 11 | S2? | |
| | | | | Time x group | 2, 34 | 2.19 | 0.127 | | | | |
| | | | | Group | 2, 34 | 0.091 | 0.913 | | | | |
| | | 48hr, short term | one way T-test | relative to 50% | 16 | 1.428 | 0.174 | | | | |
| | | 48hr, long term | one way T-test | relative to 50% | 15 | 4.24 | 0.001 | ** | | | |
| | | 2 wk, short term | one way T-test | relative to 50% | 10 | 0.77 | 0.459 | | | | |
| | | 2 wk, long term | one way T-test | relative to 50% | 10 | 3.261 | 0.009 | ** | | | |
| | | 4 wk, short term | one way T-test | relative to 50% | 10 | 2.163 | 0.056 | | | | |
| | | 4 wk, long term | one way T-test | relative to 50% | 10 | 1.009 | 0.337 | | | | |
| | | Short vs long term, partner 1 | partner time | pearson, spearman | | 0.335, 0.341 | 0.043, 0.039 | * | n = 37 | 1D | |
| | | | partner huddle | pearson, spearman | | 0.347, 0.417 | 0.036, 0.010 | * | n = 37 | | |
| | | | percent phuddle | pearson, spearman | | 0.290, 0.387 | 0.081, 0.018 | * | n = 37 | | |
| | | | novel time | pearson, spearman | | 0.413, 0.423 | 0.011, 0.009 | * | n = 37 | | |
| | | | novel huddle | pearson, spearman | | 0.294, 0.353 | 0.077, 0.032 | * | n = 37 | | |
| | | Ndist/Pdist corr with PctPhuddle | Short term | pearson, spearman | | 0.782, 0.827 | 6.7e-9, 1.55e-10 | *** | n = 38 | | |
| | | Ndist/Pdist corr with PctPhuddle | Long term | pearson, spearman | | 0.760, 0.790 | 1.55e-10, 4.73e-8 | *** | n = 37 | | |
| Huddle time | 1 | Partner huddle time | paired t-test | Time (RM) | 36.000 | 0.192 | 0.849 | | n = 37 | 1B | no data for 2242, long term PPT |
| Huddle time | 1 | Novel huddle time | paired t-test | Time (RM) | 36.000 | 2.347 | 0.025 | * | n = 37 | 1B | |
| Avg. distance from tethered partner | 1 | Distance when in chamber | RM-ANOVA | Time (RM) | 1, 36 | 4.090 | 0.051 | | n = 37 | 1C | no data for 2242, long term PPT |
| | | | | Tethered animal (RM) | | 24.252 | 0.000 | *** | | | |
| | | | | Time x tethered | | 1.534 | 0.224 | | | | |
| | | | 48 hr; Paired t-test short term | | 14 | -3.899 | 0.002 | ** | n = 15 | | |
| | | | 48 hr; Paired t-test long term | | 14 | -3.727 | 0.002 | ** | | | |
| | | | 2 wk; Paired t-test short term | | 10 | -2.765 | 0.020 | * | n = 11 | | |
| | | | 2 wk; Paired t-test long term | | 10 | -3.615 | 0.005 | *** | | | |
| | | | 4 wk; Paired t-test short term | | 10 | -3.367 | 0.007 | *** | n = 11 | | |
| | | | 4 wk; Paired t-test long term | | 10 | -2.334 | 0.042 | * | | | |
| Distance from tethered | 1 | PminusN distance | RM-ANOVA | Time (RM) | 1, 36 | 1.534 | 0.224 | | n = 37 | not shown | no data for 2242, long term PPT |
| | | | RM-ANOVA | Time (RM) | 1, 34 | 1.224 | 0.276 | | | | |
| | | | | Group | 2, 34 | 0.545 | 0.585 | | | | |
| | | | | Time x group | 2, 34 | 1.276 | 0.292 | | | | |
| Total distance | 1 | Total distance traveled by group | RM-ANOVA | Time (RM) | 1, 34 | 13.993 | 0.001 | ** | | | |
| | | | | Group | 2, 34 | 3.031 | 0.061 | | | | |
| | | | | Time x group | 2, 34 | 1.159 | 0.326 | | | | |
| | | Total distance | short term vs long term | pearson, spearman | | 0.370, 0.284 | 0.026, 0.093 | ** | | | |
| Huddle time | 2 | 48hr, 2nd partner; Partner huddle time | RM-ANOVA | Time (RM) | 1, 12 | 3.392 | 0.090 | | n = 13 | 2A | |
| Huddle time | 2 | 48hr, 2nd partner; Novel huddle time | RM-ANOVA | Time (RM) | 1, 12 | 2.106 | 0.172 | | n = 13 | | |
| Huddle time | 2 | 2wk, 2nd partner; Partner huddle time | RM-ANOVA | Time (RM) | 1, 9 | 2.754 | 0.131 | | n = 10 | 2B | |
| Huddle time | 2 | 2wk, 2nd partner; Novel huddle time | RM-ANOVA | Time (RM) | 1, 9 | 6.424 | 0.032 | * | n = 10 | | |
| Huddle time | 2 | 4wk, 2nd partner; Partner huddle time | RM-ANOVA | Time (RM) | 1, 10 | 0.322 | 0.577 | | n = 11 | 2C | |
| Huddle time | 2 | 4wk, 2nd partner; Novel huddle time | RM-ANOVA | Time (RM) | 1, 10 | 0.343 | 0.571 | | n = 11 | | |
| Avg. distance from tethered partner | 2 | 48hr: Distance when in chamber | RM-ANOVA | Time (RM) | 1, 12 | 6.405 | 0.026 | * | n = 13 - 15 | 2D | excludes 2262, 2267 from long-term |
| | | | | Tethered animal (RM) | | 5.007 | 0.045 | * | | | |
| | | | | Time x tethered | | 1.669 | 0.221 | | | | |
| | | | Paired t-test short term | | 14 | -3.727 | 0.002 | ** | | | |
| | | | Paired t-test long term | | 12 | -1.371 | 0.196 | | | | |
| Avg. distance from tethered partner | 2 | 2wk: Distance when in chamber | RM-ANOVA | Time (RM) | 1, 9 | 2.636 | 0.139 | | n = 10 - 11 | 2E | excludes 1872 |
| | | | | Tethered animal (RM) | | 8.734 | 0.016 | * | | | |
| | | | | Time x tethered | | 3.290 | 0.103 | | | | |
| | | | Paired t-test short term | | 10 | -3.615 | 0.005 | ** | | | |
| | | | Paired t-test long term | | 9 | -0.623 | 0.549 | | | | |

| | | | | | | | | | | | |
|---|---|---|---|---|---|---|---|---|---|---|---|
| Avg. distance from tethered partner | 2 | 4 wk: Distance when in chamber | RM-ANOVA | Time (RM) | 1, 10 | 1.655 | 0.227 | | n = 11 | 2F | excludes 2242 |
| | | | | Tethered animal (RM) | | 19.06 | 0.001 | | | | |
| | | | | Time x tethered | | 1.872 | 0.201 | | | | |
| | | | Paired t-test short term | | 10 | -2.334 | 0.042 | * | | | |
| | | | Paired t-test long term | | 10 | -4.1 | 0.002 | ** | | | |
| Partner preference | 2 | Proportion partner huddle time | RM-ANOVA | Time (RM) | 1, 31 | 4.991 | 0.033 | * | | 2GHI (maybe combined) | excludes 1872, 2242, 2262, 2267 |
| | | | | Time x group | 2, 31 | 3.263 | 0.052 | | | | |
| | | | | Group | 2, 31 | 0.342 | 0.712 | | | | |
| | | 48hr, short term | one way T-test | relative to 50% | 14 | 2.584 | 0.022 | * | n = 15 | | |
| | | 48hr, long term | one way T-test | relative to 50% | 12 | 1.119 | 0.285 | | n = 13 | | |
| | | 48hr | Paired t-test short vs long term | Percent huddle | 12 | 2.031 | 0.065 | | | | excludes 1872 |
| | | 2 wk, short term | one way T-test | relative to 50% | 10 | 3.833 | 0.003 | ** | n = 11 | | |
| | | 2wk, long term | one way T-test | relative to 50% | 9 | 0.194 | 0.850 | | n = 10 | | |
| | | 2 wk | Paired t-test short vs long term | Percent huddle | 9 | 3.297 | 0.009 | ** | | | |
| | | 4wk, short term | one way T-test | relative to 50% | 10 | 1.985 | 0.075 | | n = 11 | | |
| | | 4 wk, long term | one way T-test | relative to 50% | 10 | 2.901 | 0.016 | * | | | |
| | | 4 wk | Paired t-test short vs long term | Percent huddle | 10 | -0.553 | 0.592 | | | | |
| Partner preference | 2 | Short vs long term, partner 2 | partner time | pearson, spearman | | 0.557, 0.593 | 0.001, 0.0003 | ** | n = 33 | excludes 1872, 2262, 2267, 2245, no data for 1850, 1867, 1924 | |
| | | | partner huddle | pearson, spearman | | 0.660, 0.732 | 0.00003, 0.000001 | ** | | | |
| | | | percent phuddle | pearson, spearman | | 0.535, 0.560 | 0.001, 0.001 | ** | | | |
| | | | novel time | pearson, spearman | | 0.594, 0.520 | 0.000268, 0.002 | ** | | | |
| | | | novel huddle | pearson, spearman | | 0.614, 0.374 | 0.000147, 0.032 | ** | | | |
| Total distance | 2 | Total distance traveled by group | RM-ANOVA | Time (RM) | 1, 30 | 1.209 | 0.280 | | n = 33 | Not shown | 4-week separated group traveled less than 2 week or 48 hour; excludes 1872, 2262, 2267, 2245, no data for 1850, 1867, 1924 |
| | | | | Group | 2, 30 | 10.252 | 0.000 | *** | | | |
| | | | | Time x group | 2, 30 | 0.212 | 0.810 | | | | |
| | | Total distance | short term vs long term | pearson, spearman | | 0.215, 0.331 | 0.229, 0.060 | | | | |
| Huddle time | 1 vs 2 | Partner 2 huddle time | ANOVA | Group | 1, 30 | 1.06 | 0.359 | | 48 hr = 14, 2 wk = 10, 4 wk = 9 | 3A | |
| | | Parnter 1 huddle time | ANOVA | Group | 1, 30 | 2.455 | 0.103 | | | | |
| Partner preference | 1 vs 2 | 48 hr group | one way T-test | relative to 50% | 13 | -0.025 | 0.981 | | n = 14 | TB 3A | |
| | | 2wk group | one way T-test | relative to 50% | 9 | 0.871 | 0.406 | | n = 10 | | |
| | | 4wk group | one way T-test | relative to 50% | 8 | 10.658 | 0.000 | *** | n = 9 | | |
| Avg. distance from tethered partner | 1 vs 2 | Distance when in chamber | RM-ANOVA | Group | 2, 30 | 2.281 | 0.120 | | 48 hr = 14, 2 wk = 10, 4 wk = 9 | 3B | excluded 2245, no data for 1850, 1867, 1942, 2242 |
| | | | | Tethered animal (RM) | 1, 30 | 17.062 | 0.000 | *** | | | |
| | | | | Group x tethered | 2, 30 | 6.480 | 0.005 | ** | | | |
| | | | 48 hr: Paired t-test | | 13 | -0.578 | 0.573 | | | | |
| | | | 2 wk: Paired t-test | | 9 | -1.064 | 0.315 | | | | |
| | | | 4 wk: Paired t-test | | 8 | -7.953 | 0.000 | *** | | | |
| Total Distance by group | 1 vs 2 | Total distance by group | ANOVA | Group | 2, 30 | 0.002 | 0.998 | | 48 hr = 14, 2 wk = 10, 4 wk = 9 | Not shown | excluded 2245, no data for 1850, 1867, 1942, 2242 |
| Mating latency 1st partner - survival analysis | 48 hr vs 2 wk vs 4wk | Latency to mate | Kaplan Meyer with Log Rank for overall comparison | | Chi sq = 1.367 | df = 2 | 0.505 | | | S1a | |
| Mating latency 2nd partner - survival analysis | 48 hr vs 2 wk vs 4wk | Latency to mate | Kaplan Meyer with Log Rank for overall comparison | | Chi sq = 0.852 | df = 2 | 0.653 | | | S1b | excludes 2242 |
| Mating latency 1st vs 2nd partner | 1st vs 2nd | Latency to mate | Kaplan Meyer with Log Rank for overall comparison | | Chi sq = 0.565 | df = 1 | 0.452 | | | | excludes 2242 |
| Proportion mated within 3 hours | 1st vs 2nd | | Fisher exact | | F = 0.313 | | 0.187 | | | | |
| Mating latency 1st vs 2nd partner | Correlation | Latency to mate | | pearson, spearman | | 0.193, 0.176 | 0.253, 0.296 | | | | non-maters assigned 10800; 9 animals (24%) did not mate early with either partner |
| Avg. distance from tethered partner for early and late mating animals | 1 | Distance when in chamber | RM-ANOVA | Time (RM) | 1, 35 | 0.220 | 0.642 | | Early mating = 23; Late mating = 14 | S1c | |
| | | | | Time x latency | 1, 35 | 0.683 | 0.414 | | | | |
| | | | | Latency (early vs non-early) | 1, 35 | 0.029 | 0.866 | | | | |
| Avg. distance from tethered novel for early and late mating [illegible] | 1 | Distance when in [illegible] | RM-ANOVA | Time (RM) | 1, 35 | 3.818 | 0.059 | | Early mating = 23; [illegible] | S1c | |
| | | | | Time x latency | 1, 35 | 0.007 | 0.933 | | | | |

| | | | | | | | | | | | |
|---|---|---|---|---|---|---|---|---|---|---|---|
| novel for early and late mating animals | [illegible] | chamber | [illegible] | Latency (early vs non-early) | 1, 35 | 0.507 | 0.481 | | Late mating = 14 | [illegible] | |
| Avg. distance from tethered partner minus novel for early and late mating animals | 1 | Distance when in chamber | RM-ANOVA | Time (RM) | 1, 35 | 1.122 | 0.297 | | Early mating = 23; Late mating = 14 | Not shown | |
| | | | | Time x latency | 1, 35 | 0.287 | 0.595 | | | | |
| | | | | Latency (early vs non-early) | 1, 35 | 0.097 | 0.757 | | | | |
| Partner preference | 1 | Proportion partner huddle time | RM-ANOVA | Time (RM) | 1, 35 | 1.21 | 0.279 | | Early mating = 23; Late mating = 14 | S1d | 2242 only had data for short term |
| | | | | Time x latency | 1, 35 | 0.458 | 0.503 | | | | |
| | | | | Latency (early vs non-early) | 1, 35 | 0.002 | 0.969 | | | | |
| | | Early maters, short term | one way T-test | relative to 50% | 22 | 1.752 | 0.094 | | | | |
| | | Late maters, short term | one way T-test | relative to 50% | 14 | 1.766 | 0.099 | | | | |
| | | Early maters, long term | one way T-test | relative to 50% | 22 | 4.13 | 0.000 | ** | | | |
| | | Late maters, long term | one way T-test | relative to 50% | 13 | 2.291 | 0.039 | * | | | |
| Partner preference | 2 | Proportion partner huddle time | RM-ANOVA | Time (RM) | 1, 32 | 4.533 | 0.041 | * | Early = 16, Late = 18 | | Excluded 1872, 2262, 2267 from long term test |
| | | | | Time x latency | | 1.296 | 0.263 | | | | |
| | | | | Latency (early vs non-early) | | 9.077 | 0.005 | ** | | | |
| | | Early maters, short term | one way T-test | relative to 50% | 17 | 10.302 | 0.998 | **** | | | |
| | | Late maters, short term | one way T-test | relative to 50% | 18 | 1.048 | 0.309 | | | | |
| | | Early maters, long term | one way T-test | relative to 50% | 15 | 3.832 | 0.002 | ** | | | |
| | | Late maters, long term | one way T-test | relative to 50% | 17 | 0.501 | 0.623 | | | | |

**Supplemental Table 2 Excluded Animals.**

| Animal | Separation Condition | Test Excluded From | Reason for Exclusion |
|---|---|---|---|
| 1872 | 2 Week | P2 Long term | Death of Partner |
| 1850 | 4 Week | P1 vs P2 | Death of Partner |
| 1867 | 4 Week | P1 vs P2 | Death of Partner |
| 1924 | 2 Week | P1 vs P2 | Death of Partner |
| 2242 | 48 Hour | P1 long term, P2 all | Death of Partner |
| 2245 | 48 Hour | P1 vs P2 | Escaped Apparatus |
| 2262 | 48 Hour | P2 long term | Camera shift so box was out of frame |
| 2267 | 48 Hour | P2 long term | Camera shift so box was out of frame |

P1 = partner 1, P2 = partner 2
